# Using Bayesian multilevel whole-genome regression models for partial pooling of estimation sets in genomic prediction

**DOI:** 10.1101/012971

**Authors:** Frank Technow, L. Radu Totir

## Abstract

Estimation set size is an important determinant of genomic prediction accuracy. Plant breeding programs are characterized by a high degree of structuring, particularly into populations. This hampers establishment of large estimation sets for each population. Pooling populations increases estimation set size but ignores unique genetic characteristics of each. A possible solution is partial pooling with multilevel models, which allows estimating population specific marker effects while still leveraging information across populations. We developed a Bayesian multilevel whole-genome regression model and compared its performance to that of the popular BayesA model applied to each population separately (no pooling) and to the joined data set (complete pooling). As example we analyzed a wide array of traits from the nested association mapping maize population. There we show that for small population sizes (e.g., < 50), partial pooling increased prediction accuracy over no or complete pooling for populations represented in the estimation set. No pooling was superior however when populations were large. In another example data set of interconnected biparental maize populations either partial or complete pooling were superior, depending on the trait. A simulation showed that no pooling is superior when differences in genetic effects among populations are large and partial pooling when they are intermediate. With small differences, partial and complete pooling achieved equally high accuracy. For prediction of new populations, partial and complete pooling had very similar accuracy in all cases. We conclude that partial pooling with multilevel models can maximize the potential of pooling by making optimal use of information in pooled estimation sets.

## INTRODUCTION

Genomic selection (Meuwissen *et al*. 2001) in animal and plant breeding rests on the accurate prediction of genomic breeding values (GEBV). An important determinant of prediction accuracy is the size of the estimation set (Daetwyler *et al*. 2010). In animal breeding, assembling large estimation sets is relatively straight forward for large dairy breeds like Holstein Friesian, where genomic selection is applied most successfully to date (Hayes *et al*. 2009). For smaller dairy cattle breeds and in particular for beef cattle breeds, however, assembling sufficiently large estimation sets within each breed is often not possible (Weber *et al*. 2012). Creation of multi-population estimation sets by pooling several breeds is therefore of great interest and subject of current research (Lund *et al*. 2014).

A similar situation exists in plant breeding, which is characterized by a high degree of structuring (Albrecht *et al*. 2014). This structuring results from the importance of keeping distinct heterotic groups for maximum exploitation of heterosis (Melchinger and Gumber 1998), from the predominance of distinct biparental populations (Riedelsheimer *et al*. 2013) and the need for specialized breeding programs targeting specific traits or environments (Windhausen *et al*. 2012). This requires that the phenotyping and genotyping resources available to a breeding program have to be allocated to multiple populations, which prevents the creation of sufficiently large estimation sets for each population. Several studies therefore investigated the merit of pooled estimation sets combining populations (Asoro *et al*. 2011; Heffner *et al*. 2011; Lorenz *et al*. 2012; Riedelsheimer *et al*. 2013; Lehermeier *et al*. 2014) or even heterotic groups (Technow *et al*. 2013; Lehermeier *et al*. 2014).

However, pooling estimation sets is complicated by genetic differences among populations, such as in linkage disequilibrium, allele frequencies or relationship structure (Windhausen *et al*. 2012; Weber *et al*. 2012; Riedelsheimer *et al*. 2013; Technow *et al*. 2014). This might be the reason why using pooled estimation sets failed to increase prediction accuracy in some applications in plant (Desta and Ortiz 2014) and animal breeding (Lund *et al*. 2014).

Therefore, Brøndum *et al*. (2012) proposed to use separate estimation sets for each population but to derive genome position specific priors from estimation results in the other population. In this way, unique genome properties of each population could be accounted for while still using information from other populations. A similar, but perhaps more formal approach is “partial pooling”, facilitated by Bayesian multilevel models (Gelman and Hill 2006; Gelman and Pardoe 2006; Gelman 2006a). In multilevel models, parameters (e.g., marker effects) are estimated specific for each population but are “shrunken” towards an overall marker effect. Both the specific and overall marker effects are estimated simultaneously from the data, thereby allowing that the former are still informed by data from the other populations. Partial pooling thus strikes a middle ground between “no pooling” (specific marker effects estimated from data of specific population only) and “complete pooling” (unspecific marker effects estimated from pooled estimation set).

Our objectives were to (i) demonstrate the use of Bayesian multilevel whole-genome regression models for genomic prediction and (ii) determine in which scenarios partial pooling might be superior over no or complete pooling of estimation sets. Our investigations were based on two publicly available maize breeding data sets and supported by a simulation study.

## MATERIALS AND METHODS

### Multilevel whole genome regression model

The model fitted to the data was

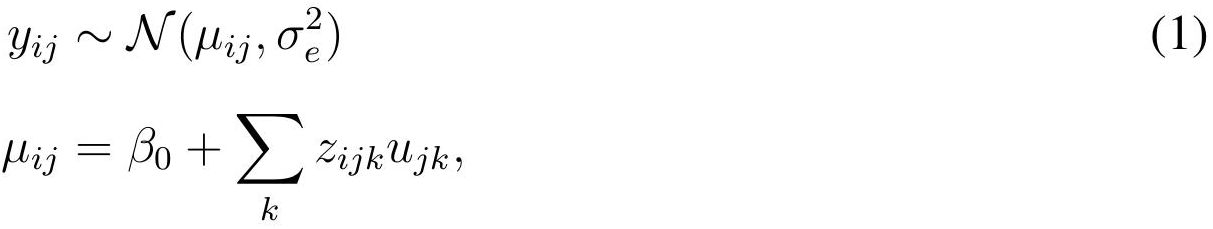

where *y*_*ij*_ was the observed phenotypic value of the *i*^*th*^ individual from the *j*^*th*^ population and *μ*_*ij*_ its linear predictor. The phenotypic data *y*_*ij*_ was centered to mean zero and scaled to unit variance. The Normal density function, which was used as likelihood, was denoted as 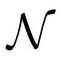 with 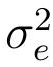 denoting the residual variance component. The common intercept was *α*_0_. Finally, *u_kj_* denoted the additive effect of the *k*^*th*^ biallelic single nucleotide polymorphism (SNP) marker in population *j*. The genotype of individual *i* from population *j* at marker *k* was represented by *z*_*ijk*_, which was the number of reference alleles, centered by twice the reference allele frequency. Which of the alleles was chosen as reference allele depended on the data set and is described below. Effects *u*_*kj*_ were only estimated when the corresponding marker was polymorphic in population *j*. Otherwise it was set to 0 and treated as a constant.

**FIGURE 1:**
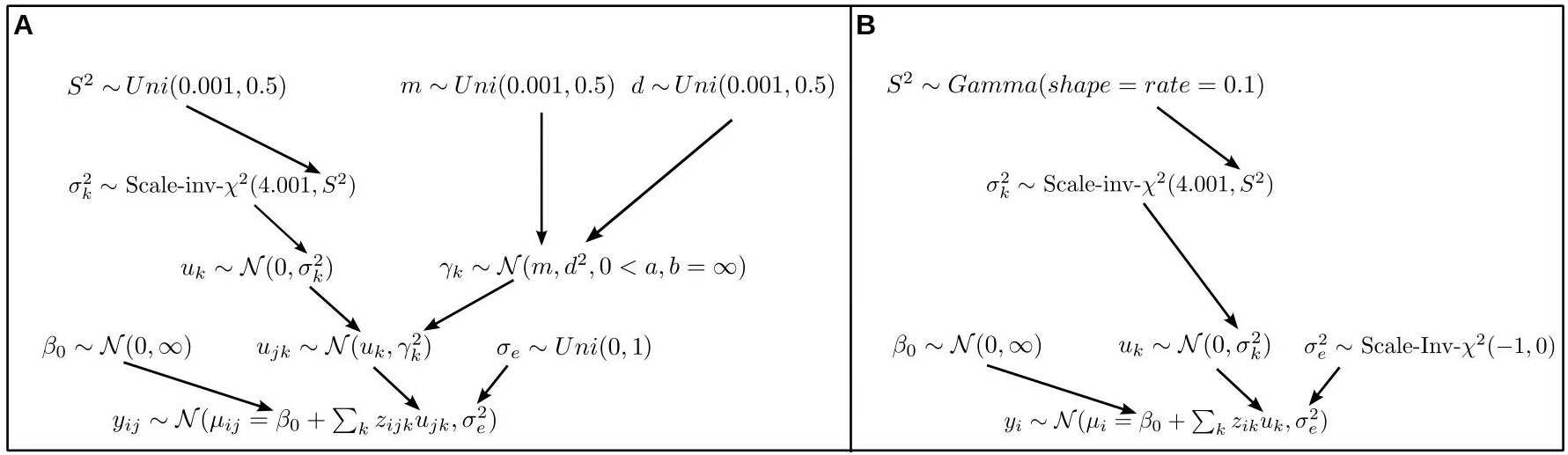
Graphical visualization of the multilevel model (A) and the conventional BayesA model (B).

The hierarchical prior distribution setup will be explained next. A graphical display is shown in Figure 1A. The prior of *u*_*kj*_ was

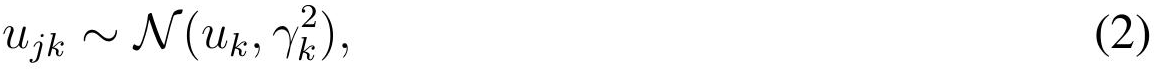

where *u*_*k*_ was the overall effect of the *k*^*th*^ marker and variance parameter 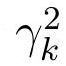 quantified the deviations of the specific effects *u*_*kj*_ from *u*_*k*_. Note that all else equal, the shrinkage toward *u*_*k*_ is the stronger the smaller 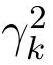.

Both parameters were associated with prior distributions themselves and estimated from the data. For *u*_*k*_ this was 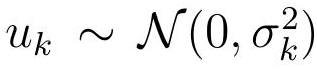. Here, the variance parameter 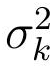 controls the amount of shrinkage towards 0. It was associated with a scaled inverse Chi-square prior with 4.001 degree of freedom and scale parameter *S*^2^. The prior for *u*_*k*_ thus corresponded to the well known “BayesA” prior (Meuwissen *et al*. 2001).

For the variance parameter 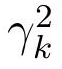, we specified

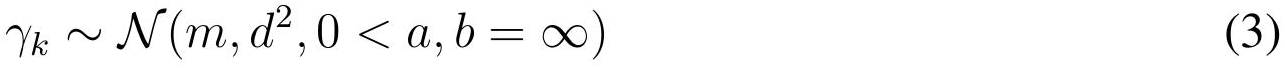

which is a Normal distribution prior on *γ*_*k*_ with mean parameter *m* and standard deviation *d*, left truncated at zero. Note that the mean of the truncated distribution 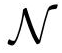 (*m*, *d*^2^, 0 < *a*, *b* = ∞), which is a function of *m*, *d* and the truncation points, can be interpreted as the “typical” deviation of the specific marker effects *u*_*kj*_ from *u*_*k*_. Higher values of this mean indicate larger deviations and viceverse. This parameter might therefore be used to quantify population divergence.

An uniform prior Uni(0:001; 0:5) was used for the hyperparameters *S*^2^, *m* and *d*. The prior for the intercept *β*_0_ was a Normal distribution with mean 0 and a very large variance. For the residual variance 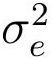 we specified a uniform distribution prior over the interval [0, 1] on *σ*_*e*_, which agrees with recommendations for uninformative priors on variance components (Gelman 2006b).

Samples from the posterior distribution were drawn with Gibbs sampling, implemented in the JAGS Gibbs sampling environment (Plummer 2003). The total number of samples was 1000, drawn from a single chain with burn in of 10000 and thinning intervals of 500. These settings ensured convergence and an effective sample size (ESS) of > 100 for all parameters (ESS of *u*_*k*_ and *u*_*jk*_ were typically > 500).

The ESS was calculated with the R (R Core Team 2013) package CODA (Plummer *et al*. 2006), which was also used to monitor convergence using diagnostic plots.

### Conventional whole genome regression model

We used the popular Bayesian whole genome regression method “BayesA” (Meuwissen *et al*. 2001), with the modifications of Yang and Tempelman (2012) pertaining to the hyperparameter *S*^2^ (see Figure 1B for a graphical representation). The linear model was

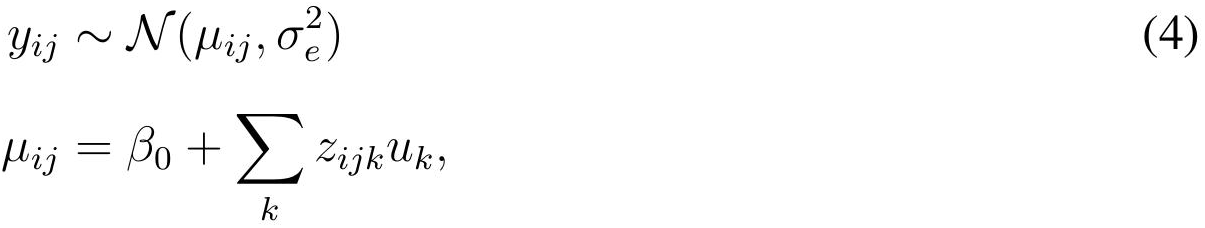

which is principally the same as in (1), with the difference that the population index *j* was dropped. For no pooling, the model was applied to each population in turn, for complete pooling to the joint data set. For 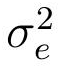 we used an improper scaled inverse Chi-square prior with -1 degrees of freedom and scale equal to zero. This is equivalent to a uniform prior on *σ*_*e*_ (Gelman 2006b), as was used for the multilevel model, but exploits conjugancy.

The BayesA Gibbs Sampler was implemented as a C routine compatible with the R statistical software environment. Again we drew a total number of 1000 samples from a single chain with burn in of 10000 and thinning of 500.

### Estimation, prediction and testing procedure

Let ∏ denote the set of *P* populations represented in the estimation set and the set of *N*_*p*_ individuals from a population in ∏ as Λ_*p*_, where *p* indexes the population in ∏. A graphical representation is presented in Figure 2. Further, let those individuals from a population in that are not in Λ_*p*_ be denoted a 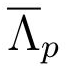 and the set of populations not in ∏ as 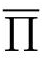 Populations in 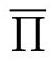 will be referred to as “new” populations. The estimation set thus comprised all individuals belonging to Λ_*p*_, for p ϵ ∏. The test set used for calculating prediction accuracy. comprised individuals in 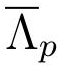 from populations in ∏ and all individuals from populations in 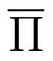 The phenotypic observations of test individuals were masked in the estimation procedure. The separation of populations into ∏ and 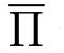 and of individuals within a population into Λ_*p*_ and 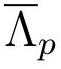 was done at random.

Within each population, prediction accuracy was computed as the correlation between GEBVs and observed phenotypic values of individuals in the testing set. The within population prediction accuracies were subsequently averaged for populations in ∏ and 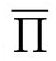 These average within population prediction accuracies will henceforth be denoted as *r*_∏_ and 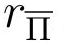 Thus, *r*_∏_ and 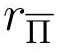 correspond to the prediction accuracy for populations represented and not represented in the estimation set, respectively.

When using partial pooling, GEBVs of individuals in 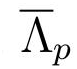 were predicted using the posterior means of the marker effects estimated for the corresponding population (*i.e*., *u*_*jk*_). GEBVs of individuals from populations in 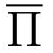 were predicted using the posterior means of the overall (unspecific) marker effects *u*_*k*_.

When using complete pooling, GEBVs of all individuals in the test set were predicted from the posterior means of marker effects *u*_*k*_ estimated from the joint data set with model (4).

Finally, when using no pooling, GEBVs of individuals in 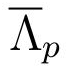 were predicted using the posterior means of the marker effects *u*_*k*_ obtained after applying model (4) to the estimation data from the corresponding set Λ_*p*_. The no pooling approach does not provide a direct way of predicting GEBVs of individuals from populations in 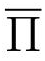 Thus,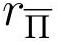 was not evaluated for the no pooling approach.

**FIGURE 2:**
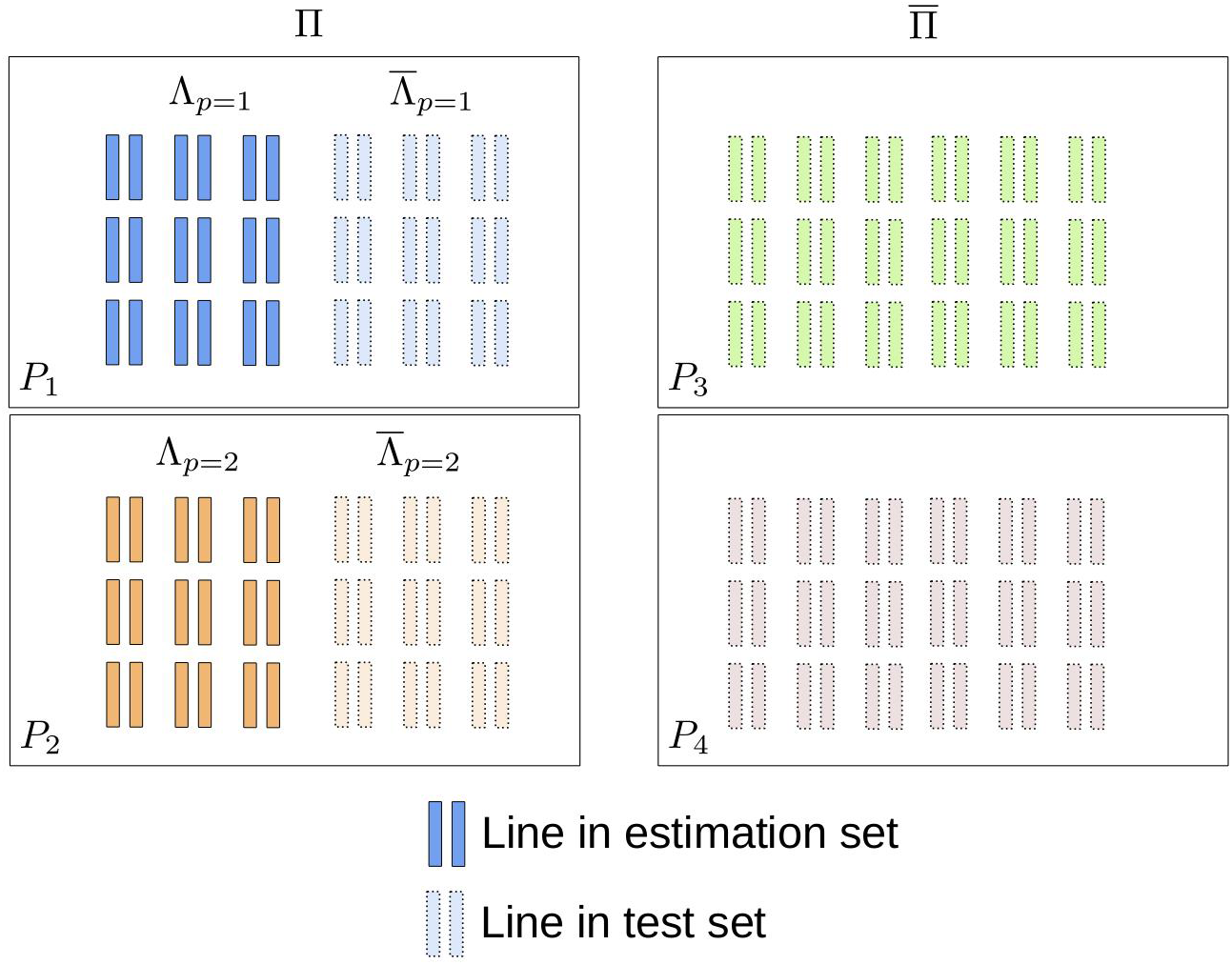
Graphical visualization of the testing strategy for evaluating prediction accuracy. The estimation set comprises Λ_1_ and Λ_2_ from populations *P*_1_ and *P*_2_ (set ∏). The prediction accuracy of lines from populations represented in estimation set (*r*_∏_) was computed from 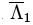 and 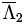, the prediction accuracy of lines from populations not represented in estimation set from lines in *P*_3_ and *P*_4_ (set ∏).

### Application to nested association mapping (NAM) maize populations

The NAM data set was obtained from http://www.panzea.org. It comprised 4699 recombinant inbred lines (RILs) from 25 biparental crosses between a genetically diverse set of maize inbred lines and line B73 as common parent (McMullen *et al*. 2009). The average population size was 188. The RILs were genotyped with 1106 polymorphic SNP markers covering the whole genome. The non-B73 allele was defined as the reference allele. We confirmed that all SNP were biallelic and thereby that the reference allele corresponded to the same nucleotide in all 25 populations. To facilitate computations, we used a thinned set of 285 markers, chosen in such a way that there was one marker per 5 cM interval, on average. A previous study showed that a density of one marker per 10 cM interval is sufficient for genomic prediction in the NAM population (Guo *et al*. 2012). We analyzed the traits days to silking (DS), ear height (EH), ear length (EL), southern leaf blight resistance (SLB), near-infrared starch measurements (NS) and upper leaf angle (ULA), which were phenotyped in multi-environment field trials. The phenotypic records used for fitting the models were averages over the single environment phenotypes. The number of environments were 10, 11, 8, 3, 7 and 9 for DS, EH, EL, SLB, NS and ULA, respectively. The traits chosen represent the major trait categories available: yield component (EL), agronomic (EH), disease resistance (SLB), flowering (DS), quality (NS) and morphology (ULA).

To investigate the effect of total number of lines *N*, number of populations *P* and number of lines per population *N*_*p*_ in the estimation set on prediction accuracy and the relative performance of the pooling approaches, the following combinations of *P* and *N*_*p*_ were considered: *P* = 5 and *N*_*p*_ = 50 and 100, *P* = 10 and *N*_*p*_ = 25, 50 and 100, *P* = 20 and *N*_*p*_ = 12.5, 25, and 50. For *P* = 20 and *N*_*p*_ = 12.5, we sampled 19 populations with 12 individuals and one with 22, which results in an average *N*_*p*_ of 12.5. The *P* and *N*_*p*_ combinations thus gave rise to *N* of either 250, 500 or 1000. For each combination of trait, *P* and *N*_*p*_, 50 estimation-testing data sets were generated by repeating the sampling of ∏ and Λ_*p*_ as described above. Throughout, the three pooling approaches were applied to the same data sets. The sampling variation between different data sets thus does not enter the comparisons among pooling approaches.

### Application to interconnected biparental (IB) maize populations

This data set was obtained from the supplement of Riedelsheimer *et al*. (2013). It comprised 635 doubled haploid (DH) lines from five biparental populations with average size of 127. The populations were derived from crosses between four European flint inbred lines. For all DH lines 16741 SNP markers polymorphic across populations were available. We replaced missing marker genotypes with twice the frequency of the reference allele, which was the allele with the lower frequency. When analyzing the data we used a thinned set of 285 markers. Because the data set did not include a map of the markers, the markers were chosen randomly.

The DH lines were phenotyped in multi-environment field trials for Giberella ear rot (GER) severity, a fungal disease caused by *Fusarium graminearum*, deoxynivalenol (DON) content (major mycotoxin produced by the fungus), ear length (EL), kernel rows (KR) and kernels per row (KpR). A more detailed description of this data set can be found in Riedelsheimer *et al*. (2013) and Martin *et al*. (2012).

As described above, populations were randomly split into Λ_*p*_ and 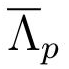 However, because there were only five populations in total, we did not exclude any populations from ∏. Set 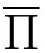 was thus empty and we did not evaluate 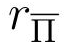

The sets Λ_*p*_ comprised 25%, 50% and 75% of the lines in each population, which corresponded to an average *N*_*p*_ of 31, 63 and 95, respectively. For each trait and percentage value of estimation individuals, 100 estimation-testing data sets generated, each time resampling the subset of 285 markers too.

### Application to simulated data set

We conducted a simulation study to specifically investigate the performance of the pooling approaches under increasing levels of differences in QTL effects among populations. The basis for the simulation were the marker genotypes of the lines in the NAM populations. To simulate genetic values, we first randomly chose 20 marker loci as QTL, which were subsequently removed from the set of observed markers. We drew additive overall effects *a*_*q*_ from a standard normal distribution. Then population specific QTL effects *a*_*jq*_ were sampled from 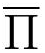. The variance parameter *τ*^2^ was chosen such that the relative standard deviation (rSD), i.e., *τ*_*q*_/*a*_*q*_, was equal to 2, 1, 0.5, 0.25 and 0.0. The greater rSD, the less similar the population specific QTL effects are. True genetic values were obtained by summing QTL effects *a*_*jq*_ according the QTL genotypes of each individual. Finally phenotypic values were simulated by adding a normally distributed noise variable to the true genetic values. The variance of the noise variable was chosen such that the heritability across populations was equal to 0.70. The average within family heritability necessarily increased with decreasing rSD, and was 0.53, 0.58, 0.64, 0.68 and 0.70 at rSD 2, 1, 0.5, 0.25 and 0.0, respectively.

Set ∏ comprised *P* = 10 populations and sets Λ_*p*_ had size *N*_*p*_ = 25. For each rSD value 50 estimation-testing data sets were generated. The QTL positions and effects were randomly generated anew for each data set. Also in this case we used a thinned set of 285 markers. Because the true genetic values were known, *r*_∏_ and 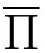 were computed as the correlation between true genetic values and GEBVs.

## RESULTS

### NAM maize populations

Trends typically held across traits. The results presented and discussed therefore apply to all traits, unless otherwise mentioned.

Increasing *N*_*p*_ while keeping *N* constant (i.e., having fewer but larger populations in the estimation set) generally increased *r*_∏_ and decreased 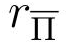 (Table 1). However, the increase in *r*_∏_ was much more pronounced than the decrease in 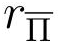.

When increasing *N*_*p*_ with constant *P* or when increasing *P* with constant *N*_*p*_, both *r*_∏_ and 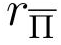 increased (Table 1). However, while in the first case, *r*_∏_ and 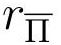 increased in similar magnitudes, the increase in *r*_∏_ was much smaller than the increase in 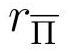 in the second case, in particular when *N*_*p*_ was high. Per definition, the accuracy of no pooling is not expected to change as long as *N*_*p*_ remains constant.

For low *P* and high *N*_*p*_, e.g., *P* = 5 and *N*_*p*_ = 100, no pooling achieved the highest *r*_∏_ and complete pooling the lowest (Table 1). For high *P* and low *N*_*p*_, e.g., *P* = 20 and *N*_*p*_ = 25, partial pooling achieved the highest *r*_∏_. Here no pooling resulted in the lowest *r*_∏_. The only exception to this was trait DS, where no pooling had a *r*_∏_ equal or higher to partial and complete pooling also for low *N*_*p*_.

Partial and complete pooling achieved virtually identical prediction accuracies 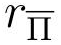 for new populations (Table 1). In general, 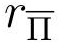 of a particular pooling approach was considerably lower than the corresponding *r*_∏_. The differences between *r*_∏_ and 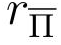 tended to be larger for high *N*_*p*_.

### IB maize populations

The prediction accuracy *r*_∏_ increased with increasing *N*_*p*_, for all traits and pooling approaches (Table 2). Averaged over traits, the increase was largest for no pooling, where the accuracy increased from an average of 0.35 at *N*_*p*_ = 31 to 0.48 at *N*_*p*_ = 95. The accuracies for the partial and complete pooling approaches increased from 0.39 and 0.38, respectively, at *N*_*p*_ = 31 to 0.48 at *N*_*p*_ = 95.

At *N*_*p*_ = 31, partial pooling had the highest *r*_∏_ for traits EL, KpR, complete pooling for traits DON and KR. For GER both had the same accuracy. The no pooling approach had the lowest *r*_∏_, except for EL and KpR, where it had the same accuracy as complete pooling. For the highest *N*_*p*_ of 95, the accuracy differences among the pooling approaches decreased. Partial pooling still had the highest accuracy for EL and KpR and the same as complete pooling for DON and GER. While never better than partial pooling, no pooling had higher prediction accuracy than complete pooling for EL and KpR.

**Table 1:**
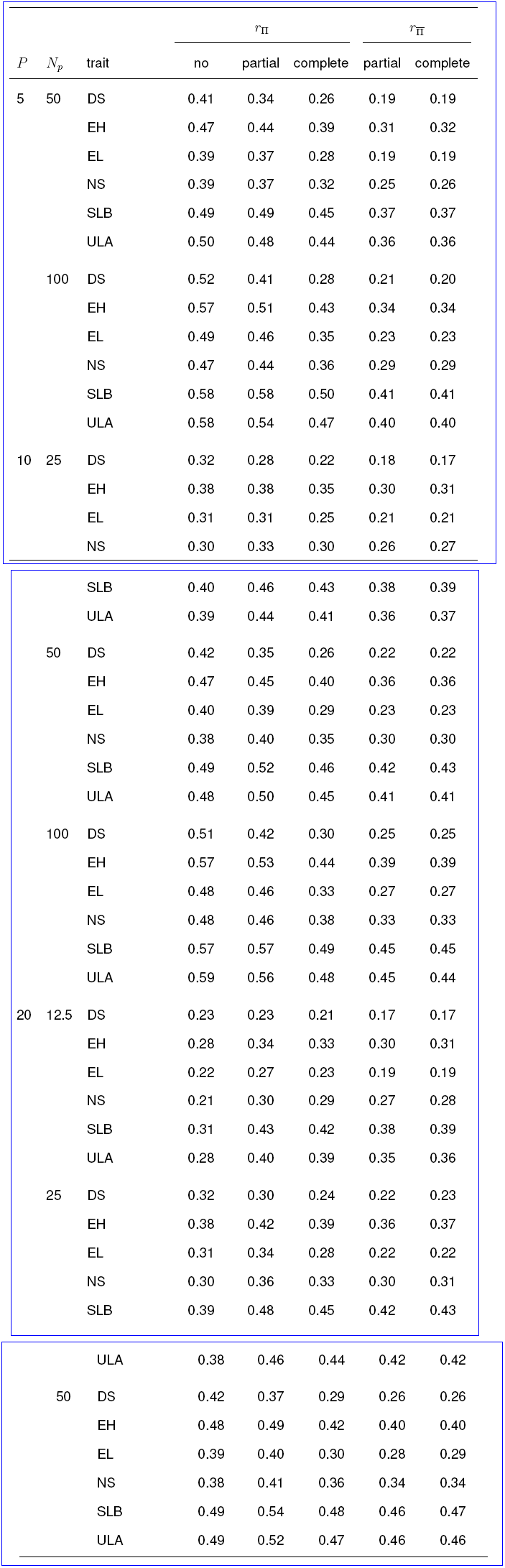
Average within population prediction accuracies in NAM maize populations

Values shown are average within population prediction accuracies for test individuals, averaged over 50 random estimation-test data splits. The standard errors were < 0.013. *P* gives the size of set ∏, i.e., the number of populations represented in the estimation set, column *N*_*p*_ gives the number of individuals from each population in ∏ that were used for estimation, i.e., the sizes of sets Λ_*p*_. The traits were: days to silking (DS), ear height (EH), ear length (EL), southern leaf blight resistance (SLB), near-infrared starch measurements (NS) and upper leaf angle (ULA).

### Simulated maize populations

For all pooling approaches, *r*_∏_ increased with decreasing rSD (Table 3). The increase for no pooling, however, was comparatively small and a result of the increasing within family heritability with decreasing rSD. The relative performance of the pooling approaches also depended on rSD. For the highest rSD value considered, no pooling had the highest r, for the intermediate rSD value of 1.0 partial pooling. For the lower rSD values complete and partial pooling achieved similarly high *r*_∏_.

Also 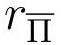 for both partial and complete pooling increased strongly with decreasing rSD and the differences to *r*_∏_ decreased (Table 3). Partial and complete pooling achieved almost identical 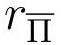.

The mean of the truncated Normal distribution prior 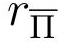 (*m*, *d*^2^, 0 < *a*, *b* = ∞) for parameter *γ*_*k*_ increased with increasing rSD. Its average values were 0.0111, 0.0153, 0.0190, 0.0269 and 0.0296 for rSD of 0.0, 0.25, 0.5, 1.0 and 2.0, respectively.

**Table 2:** Average within population prediction accuracies in interconnected biparental maize Populations

| $N_p$ | Trait | Pooling | | |
| --- | --- | --- | --- | --- |
|  |  | no | partial | complete |
| 31 | EL | 0.31 | 0.33 | 0.31 |
|  | DON | 0.38 | 0.44 | 0.46 |
|  | GER | 0.38 | 0.43 | 0.43 |
|  | KR | 0.46 | 0.50 | 0.52 |
|  | KpR | 0.21 | 0.23 | 0.21 |
| 62 | EL | 0.40 | 0.41 | 0.39 |
|  | DON | 0.47 | 0.51 | 0.51 |
|  | GER | 0.47 | 0.50 | 0.49 |
|  | KR | 0.53 | 0.56 | 0.58 |
|  | KpR | 0.28 | 0.29 | 0.27 |
| 95 | EL | 0.44 | 0.46 | 0.43 |
|  | DON | 0.51 | 0.53 | 0.53 |
|  | GER | 0.51 | 0.53 | 0.53 |
|  | KR | 0.56 | 0.58 | 0.59 |
|  | KpR | 0.31 | 0.32 | 0.30 |

Values shown are average within population prediction accuracies for test individuals, averaged over 100 random estimation-test data splits. Standard errors were < 0.01. *Np* denotes the average number of individuals per population in the estimation set. The traits were ear length (EL), deoxynivalenol content (DON), Giberella ear rot severity (GER) kernel rows (KR) and kernels per row (KpR)

**Table 3:** Average prediction accuracies for simulated maize populations

| rSD | $r_{\Pi}$ | | | $r_{\overline{\Pi}}$ | |
| --- | --- | --- | --- | --- | --- |
|  | no | partial | complete | partial | complete |
| 0.0 | 0.54 | 0.89 | 0.89 | 0.89 | 0.89 |
| 0.25 | 0.51 | 0.84 | 0.85 | 0.84 | 0.84 |
| 0.5 | 0.50 | 0.76 | 0.76 | 0.73 | 0.73 |
| 1.0 | 0.48 | 0.57 | 0.53 | 0.48 | 0.49 |
| 2.0 | 0.44 | 0.41 | 0.30 | 0.20 | 0.21 |

Values shown are average within population prediction accuracies for test individuals, averaged over 50 random estimation-test data splits. Standard errors were < 0:015. rSD is the relative standard deviation of simulated population specific QTL effects.

## DISCUSSION

### Comparison of pooling approaches

Partial pooling allows estimation of population specific marker effects while still facilitating “borrowing” of information across populations. It is therefore a compromise between no pooling, which models unique characteristics of each population but ignores shared information, and complete pooling, in which the opposite is the case.

When population sizes *N*_*p*_ are sufficiently large, borrowing information from other populations is not required for achieving high prediction accuracy of new individuals from the same population (*r*_∏_). Further enlarging estimation sets by pooling with other populations might then even be detrimental (Riedelsheimer *et al*. 2013). This explains why no pooling was the most accurate approach when *N*_*p*_ was large (e.g., >= 50), particularly in the NAM population, and why it profited most from increases in *N*_*p*_. Therefore, pooling of estimation sets is most promising if *N*_*p*_ is small due to budget of other constraints. We indeed observed that pooling was more accurate than no pooling when *N*_*p*_ was small (e.g., < 50). The superiority of either pooling approach over no pooling also increased with increasing *P*, because information from more populations was available, which is not used in no pooling. Thus, pooling is expected to most advantageous when *P* is relatively high and *N*_*p*_ low. Whether partial or complete pooling is the better approach will then also depend on the similarity of the pooled populations. The greater the similarity, the relatively better complete pooling is expected to perform, because the ability to estimate population specific marker effects becomes less important. In this situation partial pooling might even be of disadvantage, because it requires estimation of many more effects which might lead to problems associated with nonidentifiability (Gelfand and Sahu 1999). The parents of the IB populations are from the same breeding program (Riedelsheimer *et al*. 2013), whereas the non-common parents of the NAM populations were chosen to be maximally diverse and comprise temperate, tropical and specialty (sweet and popcorn) maize germplasm (McMullen *et al*. 2009). Accommodating for unique characteristics of the populations is therefore more important in NAM than in IB, which might explain why complete pooling was always inferior to partial pooling in the former but often equal or even superior in the latter and also why no pooling never achieved the highest prediction accuracy in IB, even for large *N*_*p*_.

The relative performance of the pooling approaches was very stable across traits in the NAM data set, with the exception of DS. For this trait the no pooling approach was generally superior, even at high *P* and low *N*_*p*_. Buckler *et al*. (2009) found evidence for an allelic series at the QTL identified for DS in the NAM population. Thus, while the positions of the QTL are conserved across populations, their effects differ. Possible reasons are presence of multiple alleles or QTL by genetic background interaction. In this situation, pooling of data is not expected to have an advantage over no pooling. This example also shows that decisions about whether to pool data or not have to be made on a by trait basis and should incorporate prior knowledge about genetic architecture, if available.

The dependence of the relative performance of the pooling approaches on the similarity of populations was also reinforced by the results from our simulation study. There we also observed that the mean of 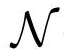 (*m*, *d*^2^, 0 < *a*, *b* = ∞), the prior distribution of 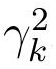, which quantifies the deviations of specific marker effects *u*_*jk*_ from the overall effect *u*_*k*_, increased with increasing simulated differences among population specific QTL effects. This was expected, but demonstrates that the data was informative for the highlevel hyperparameters. Averaged over *P* and *N*_*p*_, this mean was largest for DS and ULA in NAM (results not shown). This might reflect the noted differences between population specifuc QTL effects for DS. Trait ULA, however, did not diverge from the pattern observed for the remainder of traits and there does not seem to be any strong indication of an allelic series as in DS (reference tba). There was also no obvious relation between the mean of 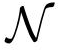 (*m*, *d*^2^, 0 < *a*, *b* = ∞) and performance of the pooling approaches in IB (results not shown).

Modeling unique characteristics of populations requires that these populations are represented in the estimation set. Prediction of individuals from new populations in 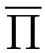 therefore has to rely on the overall, unspecific marker effects *u*_*k*_, in both partial and complete pooling. It was thus expected that both achieved very similar prediction accuracies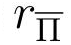 for new populations.

Our results demonstrate that partial pooling is able to model unique characteristics of populations within the estimation set without compromising on the ability of prediction of individuals from new populations. This is one reason why Gelman (2006a) see the the greatest potential of partial pooling with multilevel models in predictive applications.

We examplified the use of multilevel models for partial pooling in the context of multiple populations, a scenario of high relevance for plant (Lehermeier *et al*. 2014) and animal (Lund *et al*. 2014) breeding. However, the concept is readily applicable in a wide array of scenarios. Examples are pooling data across multiple top-cross testers or environments, as is of particular relevance in plant breeding (Albrecht *et al*. 2014). Extending the models to more than two levels is straightforward, too, for example for pooling multiple populations from multiple heterotic groups or breeding programs.

### Alternative approaches to partial pooling

There are alternatives to multilevel models for partial pooling. Brøndum *et al*. (2012) leveraged information across populations by using results obtained from one population to derive genome position specific priors for the analysis of another. For example, when there were two populations A and B, then A was analyzed first and result so obtained used as prior information when analyzing B. One disadvantage of their approach is that because analyses are done sequentially, information is not shared simultaneously among populations. In the example above, information from A is used for B but not vise verse. To use information from B for A, the analyses had to be repeated in reverse order. It is also not obvious how the approach of Brøndum *et al*. (2012) can be generalized to more than two populations or to prediction of individuals from new populations. Another potential source of concern is that the priors derived from population A are too informative to allow substantial Bayesian learning, especially when population B is small (Gelfand and Sahu 1999; Gianola 2013).

Lund *et al*. (2014) proposed to consider phenotypic observations from different populations as different traits and to analyze pooled data sets with multi-trait models. This would facilitate simultaneous sharing of information across populations through covariances. When the number of populations becomes large this might proof challenging, however, because of the need of estimating large unstructured covariance matrices. The problem is exacerbated when unique covariance matrices are estimated for each marker, as would be necessary to accommodate for varying linkage phases between markers and QTL among populations (Lund *et al*. 2014). In this case too, prediction of individuals from new populations would not be possible directly.

Schulz-Streeck *et al*. (2012) proposed a model that simultaneously fits main and population specific marker effects (*u*_*snp*_ and *u*_*psnp*_ in their notation). The principal difference to our approach is that both effects are on the same hierarchical level, such that the genetic value of an individual is modeled as the sum of *u*_*snp*_ and *u*_*psnp*_. As a consequence, both sets of marker terms “compete” for the same underlying information. This might compromise the ability of prediction in new populations which has to be based on *u*_*snp*_. Prediction targeting individuals from new populations was not attempted by the authors, however.

### Composition of estimation set

Increasing the number of individuals from a population in the estimation set (*N*_*p*_) always increased prediction accuracy for untested individuals from the same population (*r*_∏_), regardless if the estimation set was further enlarged by individuals from other populations (partial and complete pooling) or not (no pooling).

However, because plant breeding programs have to operate under budget constrains, optimum allocation of resources is of great importance for maximizing the potential of genomic selection (Lorenz 2013; Riedelsheimer and Melchinger 2013). With a fixed budget for phenotying that is proportional to *N*, the number of populations *P* and the number of individuals per population *N_p_* have to be optimized under the constraint that *N* = *PN*_*p*_. Such an optimization could be accomplished using basic theory about response to selection (Falconer and Mackay 1996) and accounting for the different prediction accuracy for populations represented and not represented in the estimation set (*r*_∏_ and 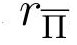, respectively), as exemplified by Technow *et al*. (2013). A key point hereby is that *r*_∏_ will increase with increasing *N*_*p*_ but it will apply to fewer populations because of the decrease in *P*. This is exacerbated by the decrease in 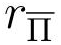 that we observed was associated with decreasing *P*. Thus, if the total number of populations is large, as is typically the case in plant breeding programs, having very low P is likely to be undesirable. In the context of plant breeding this and other studies, most recently Lehermeier *et al*. (2014), showed that pooling data across populations can at least partly compensate for low *N*_*p*_ if populations are related and there is evidence for the merit of pooling very divergent germplasm too (Technow *et al*. 2013). Using pooled estimation sets therefore has the potential to allow for high P without compromising too much on 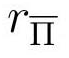 We showed that partial pooling with multilevel models can further enhance this potential by making optimal use of the information in pooled estimation sets.

